# Thiamethoxam exposure deregulates short ORF gene expression in the honey bee and compromises immune response to bacteria

**DOI:** 10.1101/853291

**Authors:** Pâmela Decio, Pinar Ustaoglu, Kamila Derecka, Ian C. W. Hardy, Thaisa C. Roat, Osmar Malaspina, Nigel Mongan, Reinhard Stöger, Matthias Soller

## Abstract

Maximizing crop yields relies on the use of agrochemicals to control insect pests. One of the most widely used classes of insecticides are neonicotinoids that interfere with signalling of the neurotransmitter acetylcholine, but these can also disrupt crop-pollination services provided by bees. Here, we analysed whether chronic low dose long-term exposure to the neonicotinoid thiamethoxam alters gene expression and alternative splicing in brains of Africanized honey bees, *Apis mellifera*, as adaptation to altered neuronal signalling. We find differentially regulated genes that show concentration-dependent responses to thiamethoxam, but no changes in alternative splicing. Most differentially expressed genes have no annotated function but encode short Open Reading Frames (sORFs), a characteristic feature of anti-microbial peptides. As this suggested that immune responses may be compromised by thiamethoxam exposure, we tested the impact of thiamethoxam on bee immunity by injecting bacteria. We show that intrinsically sub-lethal thiamethoxam exposure makes bees more vulnerable to normally non-pathogenic bacteria. Our findings imply a synergistic mechanism for the observed bee population declines that concern agriculturists, conservation ecologists and the public.

## INTRODUCTION

The western honey bee *Apis mellifera* is highly beneficial for human societies. Besides honey production, this semi-domesticated insect plays a critical role in sustaining global food security through the provision of managed pollination services that contribute to increased yields of many crops. Globally, crop productivity is also enhanced by the application of pesticides, including insecticides. Agrochemicals are, however, among the key factors implicated in contributing to declining bee health and abundance and there is a need to find a balance between the necessity of insecticide applications and the unintended effects to non-target species ^1–6^.

Compared to organophosphate pesticides, neonicotinoids are relatively safe for vertebrates and have been one of the most important classes of insecticides for the last three decades ^7^. Neonicotinoids can be applied to seeds and they then spread systemically within growing plants, killing leaf-eating pests. This reduces non-target effects compared to spraying but still retains toxic burden for pollinators as neonicotinoids continue to be found in wildflowers adjacent to treated fields ^8,9^.

Neonicotinoids act as agonists, competing with the neurotransmitter acetylcholine in binding to nicotinic acetylcholine receptors (nAChR) ^10–12^. The increased toxicity for insects is thought to be caused by the characteristically high nAChR density within insect nervous systems [reviewed in ^13^]. Detailed knowledge of cellular and molecular effects of insecticide exposure is required to mitigate negative effects or refine target specificity. Changes in gene expression and processing of RNAs, including alternative splicing, are among the mechanisms available to an organism to adapt to environmental perturbations ^14,15^. Sub-lethal exposure to xenobiotics, such as pesticides, can alter alternative splicing ^16–18^. Since many ion channel genes and other important neuronal genes undergo extensive alternative splicing, they are prime targets for changes induced by xenobiotics ^17,19^. Moreover, sub-lethal uptake of some neonicotinoids affects the honey bee brain, impairing foraging behaviour, flight, navigation, communication, learning and memory ^20–25^.

Neonicotinoid exposure has also been linked to a decline in bee health, including a reduction of immune competence ^26–28^. Insects do not have antibodies and rely on the innate immune system to fight microbial infections;; both cellular and humoral responses have been identified ^29,30^. The cellular response leading to phagocytosis or encapsulation of pathogens is mediated by three types of haematopoietic cell lineages ^29,31,32^. The humoral immune response is activated by Toll and Imd pathways [reviewed in ^33^], leading to the expression of short (10-100 amino acids) antimicrobial peptides (AMPs) which are highly diverse among insect species ^34^. The Toll-pathway is triggered by Gram-positive bacteria and fungi, ultimately leading to expression of AMPs via the NF_kappa_-B transcription factor Dif, that are then secreted from the fat body into the haemolymph ^29,35,36^. The Immune deficiency (Imd) pathway leads to expression of a different set of AMPs via the NF_kappa_-B transcription factor relish in response to peptidoglycans from primarily Gram-negative bacterial through the activation of pattern recognition receptors and a complex intracellular signalling cascade ^29,30,35^. In addition to the fatbody, AMP expression activated by the Imd pathway has also been detected in glial cells in the brain ^37–40^.

We have previously shown that worker-bee larvae in colonies contaminated with the neonicotinoid imidacloprid, have altered expression of genes belonging to lipid-carbohydrate-mitochondrial metabolic networks ^41^. We have further demonstrated that sub-lethal exposure to thiamethoxam, another neonicotinoid, can cause impairment in the midgut and brain of the Africanized *Apis mellifera*, as well as contribute to a reduction in honey bee lifespan ^9,42–44^. In this study, we analysed the effects of chronic, sub-lethal, thiamethoxam exposure on genome-wide gene expression and alternative splicing in the brains of honey bee workers. We found 52 differentially regulated genes showing a concentration dependent response to thiamethoxam exposure. Most of these genes have no annotated function but the vast majority are characterized by encoding short Open Reading Frames (sORFs), half of which are predicted to encode antimicrobial peptides. This result suggested that immune responses may be compromised by thiamethoxam exposure. Indeed, we found that thiamethoxam-exposed bees that were also infected with bacteria had greatly decreased viability compared to infected but chemically unexposed bees. Overall, our results suggest that thiamethoxam makes bees vulnerable to bacterial infection by compromising immune responses.

## MATERIALS AND METHODS

We assessed whether the neonicotinoid thiamethoxam alters gene expression in the brain of Africanized honey bees, *Apis mellifera*, after long-term exposure to field-relevant concentrations.

## Thiamethoxam exposures

Genetically unrelated bees from three unrelated hives from different locations in the apiary of the Biosciences Institute, *UNESP*, Rio Claro-SP (Brazil). Brood frames were removed from three different colonies and kept for 24 hours in biochemical oxygen demand (BOD) at 34° C and relative humidity of 80 ± 10%. The newly emerged bees (aged 0 to 24h) were marked with special non-toxic pen (Posca pen) and returned to their respective colonies until they were collected after 15 days. All experiments followed the recommendations of the guidelines for xenobiotic assessments on bees (OECD, 1998a, 1998b).

Thereafter, bees were kept in groups of 20 individuals in small cages (12 cages/treatment) within an incubator at 32° C. Bees were fed *ad libitum* with sugar solution (1:1 water and inverted sugar) from pierced 2 ml Eppendorf tubes for the control group (dataset A, Supplemental Data 1) and, for toxin exposure treatments, thiamethoxam (Sigma) was added at 2 ng/ml (low dose, LD, dataset B, Supplementary Data 1) or at 50 ng/ml (high dose, HD, dataset C, Supplementary Data 1). Sugar-feeders were re-filled before they were empty to guarantee continuous consumption. The tested concentrations were chosen based on the range of realistic field concentrations found in pollen and nectar ^9,45–48^. Assuming that caged worker bees consume about 22 µl of 50% sugar per day ^49^ the bees would have ingested 0.44 ng and 11 ng after ten days for the low and high dose exposure, respectively. After ten days, bees were cold-anaesthetised and their brains were dissected for RNA extraction.

### RNA extraction, Illumina sequencing, analysis of differential gene expression and splicing

Total RNA was extracted from ten dissected brains per sample and three replicates were done for the control, the low and the high dose treatment (dataset A, Supplemental Data 1). To extract total RNA, brains were cracked in liquid nitrogen with a pestle and then homogenized in 50 µl of Tri-reagent (SIGMA). Then the volume was increased to 500 µl and proceeded following the manufacturer’s instructions. Total RNA was then stored in 70% ethanol and transported at ambient temperature from Brazil to the UK.

Total RNA was treated with DNase I (Ambion) and stranded libraries for Illumina sequencing were prepared after poly(A) selection from total RNA (1 μg) with the TruSeq stranded mRNA kit (Illumina) using random primers for reverse transcription according to the manufacturer’s instructions. Pooled indexed libraries were sequenced on an Illumina HiSeq2500 to yield 28–41 million paired-end 125 bp reads for three control, three low dose and three high dose samples. After demultiplexing, sequence reads were aligned to the *Apis mellifera* genome (Amel-4.5-scaffolds) using Tophat2.0.6 ^50^. Differential gene expression was determined by Cufflinks-Cuffdiff and the FDR-correction for multiple tests to raw P values, with p < 0.05 considered significant ^51^. Illumina sequencing and differential gene expression analysis was carried out by Fasteris (Switzerland). Sequencing data are deposited in GEO under GSE132858 (https://www.ncbi.nlm.nih.gov/geo/query/acc.cgi?acc=GSE132858).

Alternative splicing was analysed by rMATS ^52^ and manually inspected by uploading bam files into the Integrated Genome Viewer ^53^ and comparing read frequency. Comparison of gene lists was made with Venny 2.1 (https://bioinfogp.cnb.csic.es/tools/venny/). Protein sequences from differentially expressed genes of bees were obtained from ensemble (http://metazoa.ensembl.org/Apis_mellifera/Info/Index) and blasted against *Drosophila* annotated proteins using flybase (http://flybase.org) to assign gene functions. Proteins with no assigned functions were scanned for motifs using the Interpro server (http://www.ebi.ac.uk/interpro/) ^54^. Short ORFs were analysed for antimicrobial peptide prediction using the following server: https://www.dveltri.com/ascan/v2/ascan.html ^55^.

### Reverse transcription quantitative polymerase chain reaction (RT-qPCR)

Reverse transcription was carried out with Superscript II (Invitrogen) as previously described ^56^. PCR was performed as described and PCR products were analysed on ethidium bromide stained 3% agarose gels ^19^. Neuronal genes used as reference for gene expression comparison were bee *erect wing* (*ewg, GB44653)* and *Amyloid Precursor Protein-like* (*Appl, GB48454*), based on expression analysis in *Drosophila* ^57,58^. To validate cDNAs obtained from RT, the following primers were used to amplify bee *ewg* with AM ewgG F (CCTGATGGTACCGTATCAATTATTCAAGTTG) and AM ewgH R (CCGTGTCCATCTTCTCCTGTGAGAATGATTTG), bee *Appl* with AM Appl F (GCGCGATTCCAGGAAACTGTTGCTGCTC) and AM Appl R2 (CTGCTGTCCAGCAGATGTTTGTAATGAG). Additional primers were GB45995 (GTTGCATTTTTACGCGTACAGTTACACGACAG) and GB45995 R (GGGAAATCCCCGGGAAGAGAGCAACTGGAG), GB45995 F (GTTGCATTTTTACGCGTACAGTTACACGACAG) and GB45995 R (GGGAAATCCCCGGGAAGAGAGCAACTGGAG), and GB47479 F (GGGCTATTTGCTATCTAAGTGATCCTCC) and GB47479 R (GGGTTTAGGAGTTTTCGTTTTAGCTGCTG) and PCR amplifications were done by 30 sec initial denaturation at 94° C, and then 40 cycles in total with 30 sec at 94° C, annealing at 48° C with 30 sec extension at 72° C for 2 cycles, then at 50° C with 30 sec extension at 72° C for 2 cycles and then at 52° C with and 60 sec extension at 72° C for 36 cycles with a final extension of 2 min at 72° C.

RNA quality and quantity were assessed using an Agilent 6000Nano Kit. 250 ng of DNAse-treated RNA (the same samples used for deep seq) was reversed transcribed using a SuperScript III Invitrogen kit at 55°C, following manufacturer’s instructions. Primers for quantitative real-time PCR (RT-qPCR) were designed by Primer Express software (exon/exon junctions included except for GB41813). Amplicon sizes were assessed in 15% acrylamide gel, PCR products were then sequenced and efficiency of every pair of primers was included in the analysis.

Duplicate samples of cDNA were amplified in Real Time PCR with SybrGreen chemistry under following conditions: denaturation 93°C for 30 sec, annealing 60°C for 30 sec, elongation 72°C for 30sec. *GB41923* was amplified with primers GB41923 F1 (CGCGTTGATCGTCATGATATTG) and GB41923 R1 (CTATAAGGAAATTTTGAGCCTTCGA), *GB40669* with primers GB40669 F1 (GGCCGGATATCGCTTCAAA) and GB40669 R1 (GTCTCTTTTATCTTTTCCTCGGAATTC), and *GB48969* with primers GB48969 F1 (TTGCAGCCGTAGCAAAAGGTA), GB48969 R1 (ACCGATTTGAGCACCTTGGT) and *bubblegum* (*bgm, GB51680*) with primers bgm F1 (CATGCACAAAGAGTACAAAAATTTCA) and bgm R1 (TGGTCCCAATTCTCCAGTAACA). Analysis of CT values was performed in Light Cycler480 software and normalization and differential expression was determined with the 2^-ΔΔCt^ method ^59^ by normalizing to the expression of *ewg (GB44653), Appl (GB48454)* and *actin* (*GB44311*) genes. *Actin* and *ewg* were amplified with primers ewg Fw2 (CCGCGTCTCCTACAGCTCTT) and ewg Rv2 (TGTAAAACTGCCGTAGGATATTGG) and actF1 (TTCCCATCTATCGTCGGAAGA) and actR1 (TTTGTCCCATGCCAACCAT), and *Appl* with primers AM appl F and AM appl R2, following ^41,60^.

### Bacterial infection assays

Thiamethoxam-induced alteration of anti-microbial peptide gene expression could disrupt bees’ immune response. To evaluate the effect of thiamethoxam on immunity, we adopted an assay procedure, initially developed for *Drosophila*, which assesses how efficiently injected non-pathogenic bacteria are cleared by anti-microbial peptides ^61^. To assay clearance in bees we used *Bacillus badius*, a normally non-pathogenic bacterium commonly found in the environment and *Ochrobactrum anthropi*, which are Gram-positive and Gram-negative members of the honey bee microbiome ^62,63^. *O. anthropi* were isolated from worker bee gut cultures by plating the gut content on LB agar plates incubated at 30° C.

Bacterial species were identified by colony PCR and ribosomal 16S sequencing: A colony was picked from an LB agar plate with a yellow tip and placed into 10 µl TE in a PCR tube and heated to 94° C for 5 min. The PCR mix was added adjusting the MgCl concentration to 1.5 mM. PCR was done for 20-40 cycles with 54° C annealing for 40 sec and 1 min extension at 72° C. A 490 bp fragment of the ribosomal 16S gene was amplified with primers 16S F (ACTGAGACACGGYCCAGACTCCTACGTC) and 16S R (GCGTGGACTACCAGGGTATCTAATCC) and sequenced with primer 16S Fseq (CTCCTACGGGAGGCAGCAGTRGGGTC). If sequences did not yield a single species, primers 16S F2 (GTGGACTACCAGGGTATCTAATCCTG) and 16S R2 (CCTACGGTTACCTTGTTACGACTTCAC) were used for amplification of a 733 bp fragment, which was sequenced by 16S R2seq (CCATGGTGTGACGGGCGGTGTGTAC).

Forager honey bees for infection assays were collected from colonies of the Winterbourne Garden of the University of Birmingham (UK). They were kept and injected with bacteria as we described previously ^19^. Bacteria for injections were freshly plated and grown overnight on LB plates. Then a single colony was used to inoculate a 5 ml LB in a 50 ml Falcon tube and grown overnight to saturation: 2 µl of this culture was then injected. Bacterial titres of cultures were determined at the time of injections by plating 100 µl of 10^5^, 10^6^ and 10^7^ dilutions in LB and counting the colonies the next day: this showed that 2-8 x 10^6^ *B. badius* or *O. anthropi* were typically injected. In some treatments the injected bacteria were diluted 10-fold (Fig. 3). Groups of 8 to 12 bees were exposed to combinations of bacterial and thiamethoxam doses and survival was assessed at 24 hours and 48 hours.

**Figure 1:**
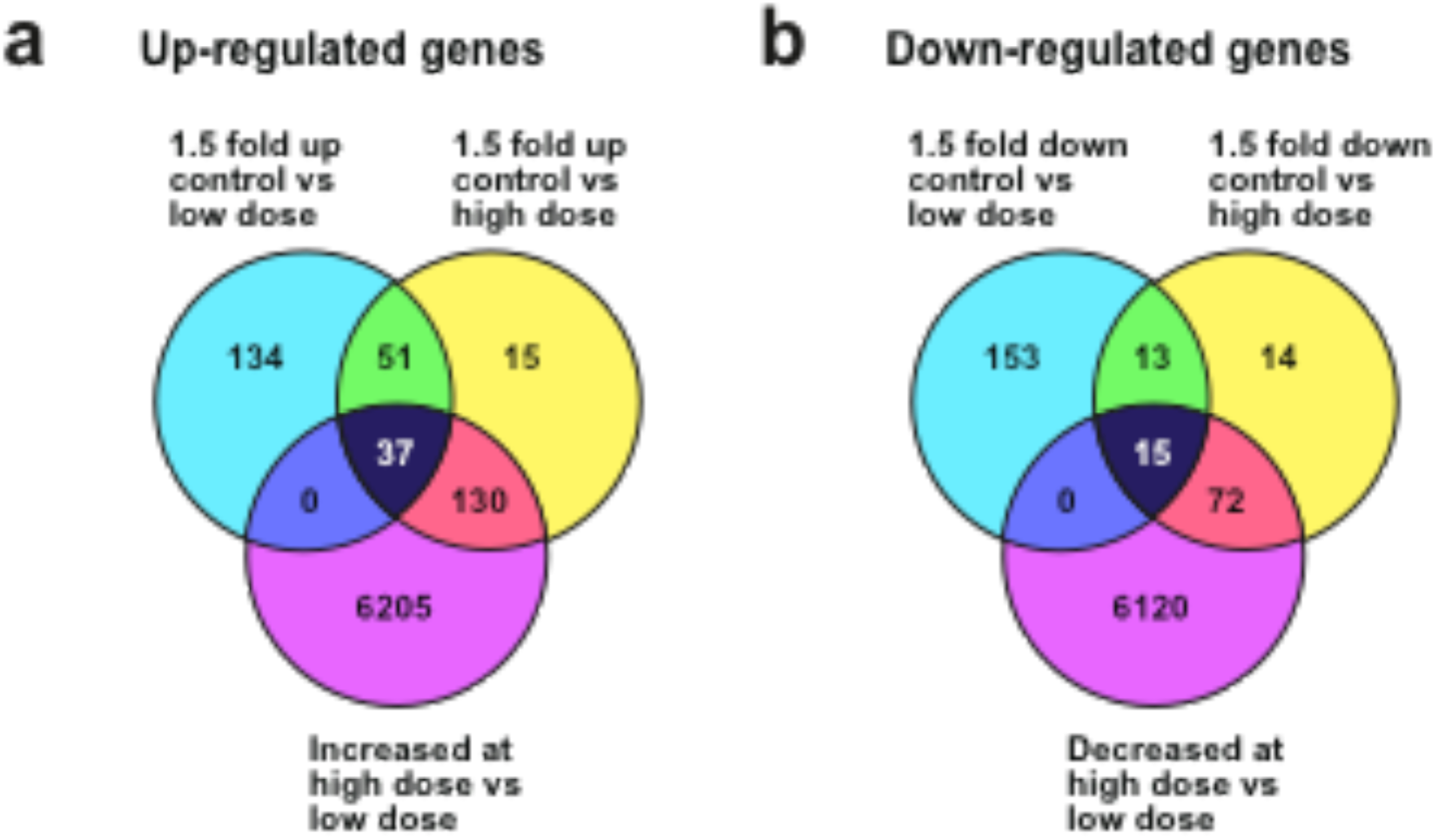
Thiamethoxam induces differential expression in a subset of genes. Venn diagrams indicating the number of differentially expressed genes between control bees and bees exposed to a low dose (left) and high dose (right) of thiamethoxam that were up-(A) or down-(B) regulated. 37 genes were up-regulated and 15 genes were down-regulated in a dose-sensitive manner.

**Figure 2:**
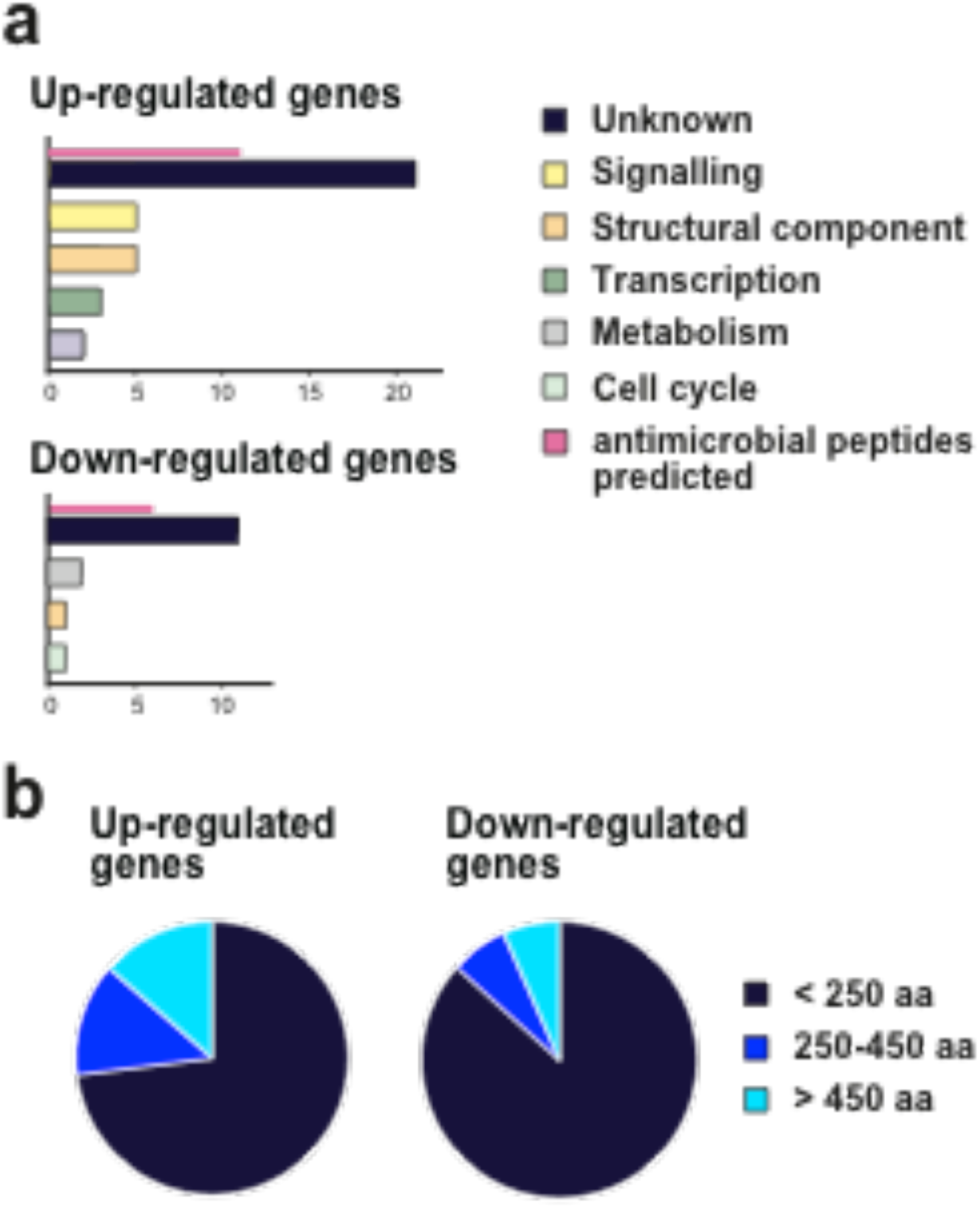
Classification of thiamethoxam induced differentially expressed genes according to function and size. (A) Numbers of genes are plotted according to functions for thiamethoxam induced differentially expressed genes with annotated functions for up-(top) and down-regulated (bottom) genes. (B) Pie charts indicating the fraction of thiamethoxam induced differentially expressed genes according the ORF length for up-(left) and down-(right) regulated genes.

**Figure 3:**
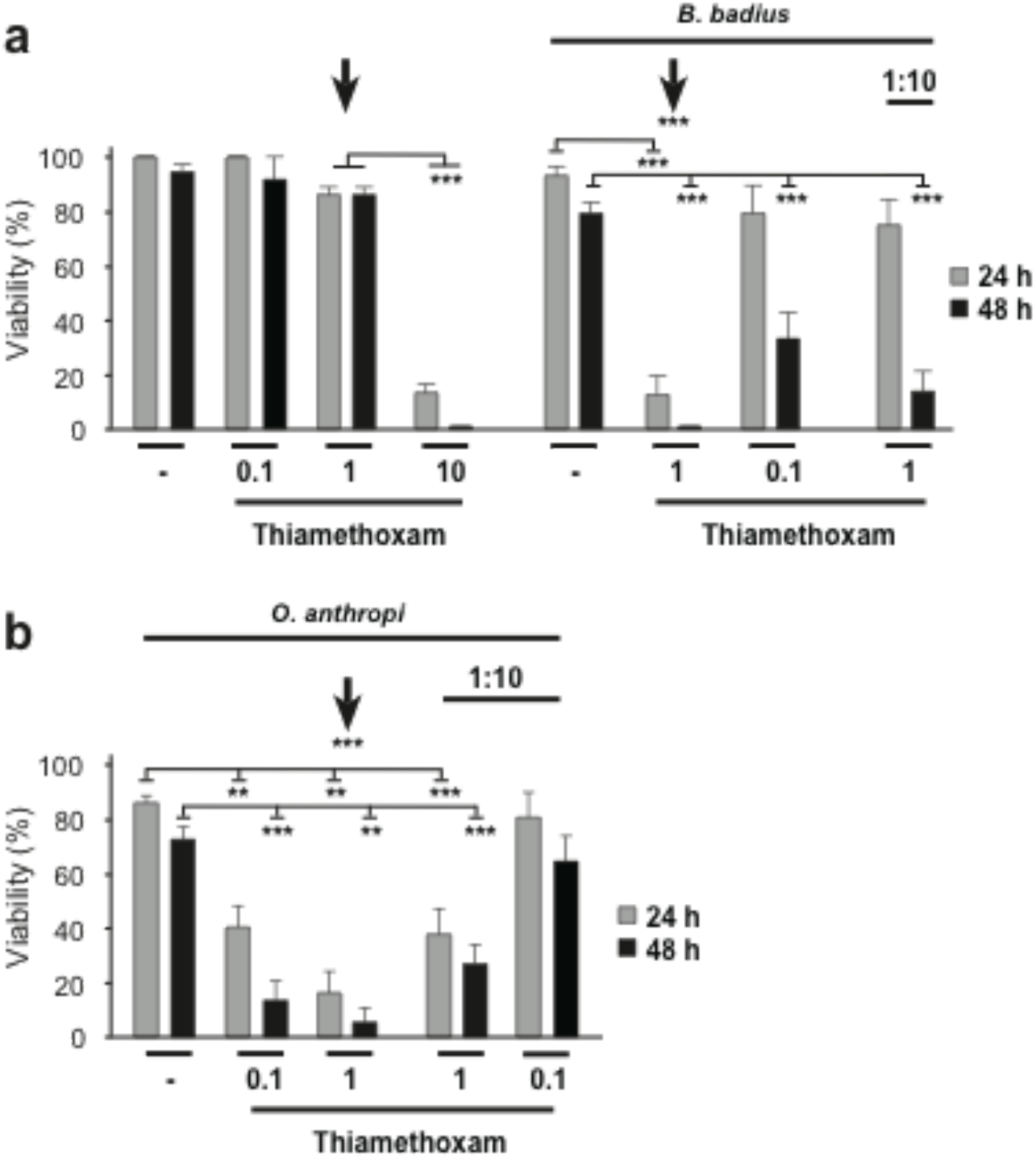
Thiamethoxam exposure makes bees vulnerable to infection by *B. badius* and *O. anthropi*. Viability of groups of 8-12 bees 24 h (grey bars) and 48 h (black bars) after injection with *B. badius* (A) or *O. anthropi* (B) (2-8 x 10^6^ and diluted 1:10) alone or together with thiamethoxam (at a range of doses, µM) shown as means and standard errors of three replicates. Arrows indicate injection of a 1 µM Thiamethoxam solution with (A right and B) or without bacteria (A left). Statistical significance is indicated by ** (p<0.01) and *** (p<0.001).

Data on survival were analysed using backwards stepwise logistic ANOVAs adopting a logit-link function and assuming quasi-binomial distributed errors. These analyses were carried out using GenStat v.19 (VSN International, Hemel Hempstead). Statistical analysis of viability were done with GraphPad prism using ANOVA followed by Tukey-Kramer post-hoc tests.

## RESULTS

### Chronic thiamethoxam exposure effects gene expression

After low (LD) and high dose (HD) exposure to thiamethoxam, there were 222 up- and 181 down-regulated genes for LD (Fig. 1a) and 233 up- and 114 down-regulated genes for HD with a 1.5 fold difference in expression compared to the control treatment (Fig. 1b;; Supplemental Table 1). From these differentially regulated genes, 37 were up-regulated and 15 were down-regulated in a dose-sensitive manner (Fig. 1;; Supplemental Table 1).

**Table 1.** Effects of thiamethoxam dose, bacterial dose and species on bee viability.

| Source | Survival after 24 hours |  |  |  | Survival after 48 hours |  |  |  |
| --- | --- | --- | --- | --- | --- | --- | --- | --- |
|  | *d.f.* | Deviance | *F* | *P* | *d.f.* | Deviance | *F* | *P* |
| *2-way logistic ANOVAs* |  |  |  |  |  |  |  |  |
| *B. badius* dose | 1 | 60.69 | 59.26 | <0.001 | 1 | 74.64 | 50.04 | <0.001 |
| Thiamethoxam dose | 1 | 73.10 | 71.38 | <0.001 | 1 | 49.92 | 33.47 | <0.001 |
| Bacteria × Thiamethoxam | 1 | 0.55 | 0.54 | 0.472 | 1 | 36.79 | 24.66 | <0.001 |
| Residual | 17 | 17.41 |  |  | 17 | 25.36 |  |  |
| Total | 20 | 121.44 |  |  | 20 | 186.72 |  |  |
| *O. anthropi* dose | 1 | 55.42 | 18.99 | <0.001 | 1 | 76.28 | 1.76 | 0.003 |
| Thiamethoxam dose | 1 | 59.99 | 20.55 | <0.001 | 1 | 44.51 | 11.12 | <0.001 |
| Bacteria × Thiamethoxam | 1 | 0.09 | 0.03 | 0.865 | 1 | 7.06 | 19.06 | 0.199 |
| Residual | 20 | 58.38 |  |  | 20 | 80.04 |  |  |
| Total | 23 | 156.51 |  |  | 23 | 191.16 |  |  |
| *3-way logistic ANOVAs* |  |  |  |  |  |  |  |  |
| Bacterial species | 1 | 15.41 |  | 0.021 | 1 | 6.21 | 0.25 | 0.622 |
| Bacterial dose | 1 | 32.67 |  | 0.002 | 1 | 14.87 | 6.28 | 0.021 |
| Thiamethoxam dose | 1 | 90.19 |  | <0.001 | 1 | 85.56 | 36.12 | <0.001 |
| Bacterial species × Bacterial dose | 1 | 3.01 |  | 0.281 | 1 | 8.73 | 3.68 | 0.069 |
| Bacterial species × Thiamethoxam | 1 | 0.13 |  | 0.204 | 1 | 6.21 | 2.62 | 0.121 |
| Bacterial dose × Thiamethoxam | 1 | 4.23 |  | 0.824 | 1 | 2.56 | 1.08 | 0.311 |
| Residual | 20 | 49.08 |  |  | 20 | 47.39 |  |  |
| Total | 26 | 155.30 |  |  | 26 | 151.08 |  |  |

To validate these results from the Illumina sequencing, we performed RT-qPCR for three of the dose-responsive genes: *GB41923*, a putative sodium-chloride co-transporter, and *GB48969, GB40669*, two genes with unknown function. We detected an expression difference for all three genes upon thiamethoxam exposure (Supplemental Fig. 1). We also validated and confirmed differential expression of *bubblegum*, encoding a very long-chain acyl-CoA synthetase, which has been found to be down-regulated in honey bee larvae exposed to the neonicotinoid imidacloprid ^41^ (Supplemental Fig. 1).

Alternative splicing has been suggested as a mechanism to adapt gene expression to environmental changes ^17,64^. We analysed the RNA-seq data for changes in alternative splicing but found no conclusive patterns (Supplementary Information 1;; Supplemental data 2).

### Dose-responsive expression occurs mostly in genes encoding uncharacterized ORFs

Next, we categorized the genes dose-responsive to thiamethoxam according to their functions, taking into account known functions of orthologues in *Drosophila* and functions deduced from annotated protein domains retrieved by BLAST analysis. Amongst the dose-sensitive genes that were up-regulated (Fig. 1A), 14 % (5/37) were assigned roles in cellular signalling (with potential links to altered neuronal function, such as olfactory and taste perception) and as structural components (cytoskeleton), and 8 % (3*/*37) were assigned functions in transcriptional regulation of gene expression (Fig. 2A). However, 59% (22*/*37) of dose-sensitive up-regulated genes and 73 % (11*/*15) of dose-sensitive down-regulated genes had neither clear orthologues in *Drosophila* nor any recognizable protein domains that would indicate a biological function (Fig. 2A), which is in contrast to about 20% of genes with unknown function in gene expression studies in *Drosophila*, ^65,66^.

### Most thiamethoxam dose-responsive genes encode short proteins

Many of the dose-responsive, differentially expressed genes with unknown function coded for short ORFs. For thiamethoxam-induced differentially up- and down-regulated genes, respectively 73 % (27*/*37) and 87 % (13*/*15) encode for genes with ORFs of 250 amino acids or shorter (Fig. 2). Using a machine learning algorithm ^55^, we predicted the 40 genes coding for peptides of ≤250 amino acids, 17 (43 %), the peptides have antimicrobial function (11/17 genes were up-regulated and 6/17 were down-regulated, Fig. 2).

### Thiamethoxam makes worker bees more vulnerable to bacterial infection

When saturated liquid cultures (2-8 Mio bacteria in 2 µl) of *B. badius* and *O. anthropi* were injected into bees that were not exposed to thiamethoxam, viability was little affected, indicating that the bee immune system usually clears the infection efficiently (Fig. 3, Table 1). In contrast, co-injection of normally sub-lethal doses of thiamethoxam together with either *B. badius* or *O. anthropi* resulted in bee death (Fig. 3, Table 1). In particular, at a dose of 1 µM thiamethoxam (arrow in Fig 3) bees generally survived for 48 h. In contrast, presence of either *B. badius* or *O. anthropi* together with thiamethoxam killed almost all bees within 48 h.

Using data on each bacterial species separately, along with replicates without bacterial exposure, showed that bee viability was negatively affected by increasing doses of thiamethoxam and of bacteria (Table 1: 2-way logistic ANOVAs). At 48 hours, a synergistic interaction between these main effects was detected when the bacterial species was *B. badius*: bee viability declined more rapidly in response to bacterial dose when bees were exposed to higher doses of thiamethoxam (Table 1, Fig. 3).

To test for differences between the effects of the two bacterial species, replicates without bacterial exposure were excluded and the effects of species, non-zero bacterial dose, thiamethoxam and their interactions were explored (Table 1: 3-way logistic ANOVAs). At 24 hours, bee viability declined with both thiamethoxam dose and bacterial dose and was lower when the bacterium applied was *O. anthropi*, without interactions (Table 1). At 48 hours, the same dosage effects were found but the species effect was not significant (Table 1). There was, however, a marginally non-significant interaction between bacterial species and bacterial dose: as probability estimates from logistic analyses are inexact, we explored the consequences of retaining this interaction in the model. This generated a further significant interaction between thiamethoxam dose and the species of bacteria applied (*F*_1,21_ = 5.85, *P*=0.025): *O. anthropi* affected bees more negatively than did *B. badius* as pesticide dose increased. We conclude that thiamethoxam suppresses the abilities of bees to cope with natural challenges of the immune system, which would normally not be fatal.

## DISCUSSION

Our study addresses the effect of thiamethoxam on gene expression at field relevant doses of thiamethoxam found in pollen and nectar. In particular we chose for these experiments a low and a high dose, which is about 0.04 ng and 1.1 ng per bee per day for a 10 day chronic long-term exposure. A number of studies have used thiamethoxam doses in the same range for analysing changes in gene expression or observing an impact on behaviour following short term exposure ^18,20–24,67,68^. In fact, it was found that chronic low-dose long-term thiamethoxam exposure altered bee activity, motor function and movement to light ^24^, but to our knowledge no previous analysis of gene expression under such conditions has been done.

A key finding of our analysis of honey bee transcriptomes is the highly enriched fraction of dose-responsive, uncharacterised genes encoding short open reading frames (sORFs). Such sORFs have only recently been recognized to encode functional peptides ^69–71^, some of which play important roles during development or have a function in the immune system ^72,73^. Intriguingly, the majority of the dose-responsive sORFs we have identified are predicted to encode peptides with antimicrobial function (anti-microbial peptides, AMPs). The sORFs we identified have not been reported in prior whole-transcriptome evaluations of neonicotinoid exposure in bee brains despite using similar concentrations of neonicotinoids ^18,67,68^. A possible explanation for the different gene set we obtained after thiamethoxan exposure is that we analysed changes after long-term, low dose exposure, while prior studies used shorter exposures. The finding that mostly sORFs, and among them many predicted AMPs are differentially expressed points to the immune system for being dysregulated by neonicotinoids. The best-characterized AMPs are expressed in the fat body upon pathogen exposure, but it is now recognized that glial cells in the brain also exert immune functions and express AMPs ^37,39,40^. Accordingly, the sORFs that we found to be differentially expressed upon chronic low dose thiamethoxam exposure might form a basal immune defence, similar to the antimicrobial environment present in saliva in vertebrates which many AMPs. Glia are support cells of neurons and are protective in a number of ways. Dysregulated AMP’s in the brain have been linked to neurodegenerative disease ^37,38,40^. Hence, the dysregulation of sORFs in response to thiamethoxam exposure could indicate disturbed communication between these two cell types beyond immune functions. Likewise, antimicrobial peptides have also been identified in having a role in learning and memory in *Drosophila*, but the molecular mechanism how AMPs enhance learning and memory remains unknown ^74^.

We noted that other studies found an overrepresentation of down-regulated genes with known function in immune and defence processes upon exposure to different types of neonicotinoids ^18,26,67,68,75^. In contrast we did not find differentially regulated immune genes including the known AMPs in the brain. AMPs are highly expressed in the fat body. For the analysis of gene expression in bee brains we dissected them from cold-anaesthetized bees and ensured that all fat body surrounding the brain was removed. It is possible, that freezing of brains prior to dissection ^67^ or residual fat body adhering to the brain contributed considerably to a different profile of immune gene expression. Immunosuppression has also observed been upon exposure to neonicotinoids co-infecting bees with viruses of *Nosema, a* unicellular parasite of bees. In particular, this affected the cellular response, but also expression of AMPs ^18,26,67,68,75^. Since we did not detect differential expression of known immune genes, the bees in our study were not infected with these known pathogens, but we cannot exclude an impact of other microbiota that are non-pathogenic in healthy bees. Moreover, our analysis focused on gene expression in the brain, where glial cells govern immune functions, which is a different humoral response than mediated by fatbody cells secreting AMPs into the hemolymph. Bees, including *Apis mellifera*, are characterised by their limited set of canonical immune genes, compared to non-social insects, such as the fruit fly *D. melanogaster* ^76–78^. Currently, only six antimicrobial-peptide genes comprising four gene families have been described in honey bees ^78^. In contrast, *Drosophila* has 20 antimicrobial-peptide genes comprising eight gene families ^29,78^. From these genes, only *defensin* is conserved between honey bees and *Drosophila*, consistent with the idea that antimicrobial peptides evolve fast to adapt to species-specific environmental conditions ^34^. Given the low number of known antimicrobial peptides in bees, it is conceivable that new (currently uncharacterised) genes encoding antimicrobial peptides are evolving. Consistent with this idea, some of the sORFs differentially regulated by thiamethoxam are bee or hymenopteran specific.

Various agrochemicals, have been shown to alter the gut microbiome ^79,80^. Our results are consistent with previous findings where neonicotinoid exposure adversely affects insect immunity ^80,81^. Specifically, we have shown that immune challenges from what are normally non-pathogenic bacteria become fatal to bees when combined with thiamethoxam exposure. How foreign bacteria enter the body of bees appears to be crucial For example, Dickel and colleagues found that oral ingestion of *Enterococcus faecalis* bacteria, increases the survival neonicotinoid-exposed, caged worker bees ^82^.In our study, the bacteria were injected between segments, which mimics punctures sustained by attacks of the *Varroa* mite. We note that pathogens can enter bee haemolymph through punctures inflicted by *Varroa destructor* mites ^63^ and thus there may be considerable mortality within hives infested with *Varroa* and also exposed to thiamethoxam.

In summary, the most prominent changes in gene expression upon long-term low-dose thiamethoxam exposure identified mostly genes that encode short ORFs, around half of which are predicted to code for antimicrobial peptides. Furthermore, thiamethoxam exposure reduced the capacity of bees to withstand microbial infection. Taken together, these findings imply that low doses of neonicotinoids may be intrinsically sub-lethal to bees but can be ultimately fatal via a weakened immune response to extrinsic pathogens. The roles of the identified genes in the immune response of bees will need to be identified to establish how bee immunity might be strengthened to resist bacterial infections.

## Authors’ contributions

P.D., P.U. and K.D. performed the experiments, T.C.R, I.C.W.H., O.M., N.M. analysed data, R.S. and M.S. supervised experiments and analysed data, R.S. and M.S. wrote the manuscript with help from P.D., P.U. and I.C.W.H.

## Acknowledgments

We thank N. Parker and the Winterbourne garden for bees, N. Parker for bee suits, V. Soller-Haussmann and K. Nallasivan for help with bee collections. G. Salmond for bacterial strains, D. Scocchia for culturing and genotyping bacteria, and L. Orsini, E. Davies and I. Haussmann for comments on the manuscript. For this work we acknowledge funding from the Foundation for Research Support of the State of São Paulo, FAPESP (2012 / 13370-8;; 2014 / 23197-7), the Biotechnology and Biological Sciences Research Council (BBSRC), the Nottingham-Birmingham Fund, and the Sukran Sinan Memory Fund.

## SUPPLEMENTAL INFORMATION AND DISCUSSION

### Analysis of alternative splicing

As alternative splicing has been suggested as a mechanism to adapt gene expression to environmental changes ^17,64^, we analysed the RNA-seq data for changes in alternative splicing. We found significant differences (Supplemental Table 2). Subsequent inspection of sequencing traces using Integrated Genome Viewer (IGV) ^53^ found that most of these genes had complex splicing patterns that showed no obvious differences in the number of sequence reads over alternatively spliced gene sections (data not shown);; therefore, we did not explore these genes further in this study.

Not to find changes in alternative splicing in response to low dose long-term exposure was unexpected, but we also did not find changes upon acute high dose xenobiotic exposure in selected genes in our earlier study on bees ^19^. In other insects, alternative splicing of the *Dscam* gene, which acts as a pattern recognition receptor in the immune response, can generate many different isoforms ^83,84^ and the splicing patterns can change upon bacterial infection ^85^. Since *Dscam* alternative splicing is robust against perturbations of splicing factors ^86,87^, the absence of changes might indicate that a specific immune challenge is required to change its splicing pattern ^85^. We previously also analysed alternative splicing in bee *elav* and *Xbp-1* genes as potential markers for defects in synapse formation and the stress response upon high dose acute thiamethoxam exposure, but also low dose chronic exposure did not affect their splicing ^19,65,66^. In contrast to most other species, bees have only a single *elav* gene making it unlikely that the lack of splicing differences is due to redundancy among close related ELAV RNA binding proteins ^88^. Potentially, alternative splicing might be changed only in a few cells in the brain, eluding detection without single cell analysis, or could in addition also be more subtle, requiring more replicates for detection of significantly altered alternative splicing changes^17^.

## Supplemental Figure legend

**Supplementary Figure 1:**
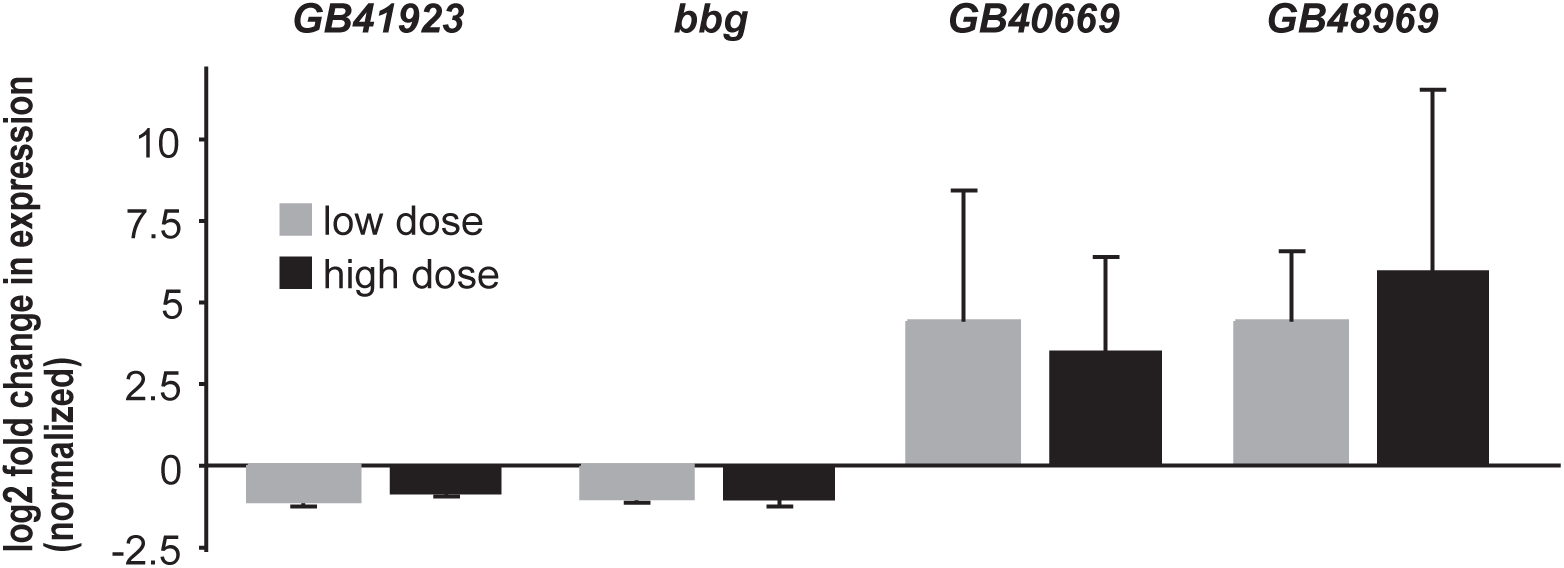
Validation of thiamethoxam induced differentially expressed genes by RT-qPCR. Means with standard error from three experiments are represented by a log2 fold change in expression levels normalized to *ewg, Appl* and *actin* genes.

